# Compensatory replacement of the BigH1 variant histone by canonical H1 supports normal embryonic development in Drosophila

**DOI:** 10.1101/789735

**Authors:** Kaili K. Li, Dongsheng Han, Fang Chen, Ruihao Li, Bing-Rui Zhou, Yawen Bai, Kai Yuan, Yikang S. Rong

## Abstract

Histone variants carry specific functions in addition to those fulfilled by their canonical counterparts. Variants of the linker Histone H1 are prevalent in vertebrates and based on the pattern of their expression, many are presumed to function during germline and the earliest zygotic stages of development. While the existence of multiple H1 variants has hampered their study in vertebrates, a single variant, BigH1, was identified in Drosophila, promising to accelerate our understanding of the biological functions of H1 and H1 variants. Here we uncovered evidence for a compensatory activity that loads maternal H1 onto BigH1-devoid chromatin. Remarkably, this H1-based chromatin state is fully functional in supporting normal embryonic development, suggesting that H1 carries the essential function of the BigH1 molecule under the same developmental context. In addition, we discovered that this compensatory replacement of BigH1 with H1 might be limited to rapidly cycling cells in early embryos.

## Introduction

Histones are the building blocks of eukaryotic chromatin. Genomic DNA wraps around a histone octamer with the linker histone H1 occupying the stretch of DNA between nucleosomes (reviewed in Fyodorov et al. 2018; and Zhou and Bai 2019). Besides these canonical histones, there are histone-like molecules called histone variants that often have specialized and essential functions (reviewed in Zink and Hake 2016, and Talbert and Henikoff 2017).

For histone H1, the existence of multiple variants is common in higher organisms (for reviews of H1 variants see Hergeth and Schneider 2015; and Pérez-Montero et al. 2016). For example, humans and mice each have eleven different H1-like proteins. The biological function of these variants has been under intense studies but not well understood. Mutational studies produced complex results in higher eukaryotes. For example, a triple knock-out of H1 and variants is needed to disrupt viability in mouse (Fan et al. 2003). However, chicken cells survive the disruption of all six annotated genes encoding H1-like proteins (Hashimoto et al. 2010), which questions the essential functions of H1 and variants at the cellular level.

Many H1 variants are present specifically in the germline and/or early development implying their roles in the corresponding developmental stages. Although biochemical characterization suggested that they maintain a specialized chromatin structure different from that based on the somatic H1 (e.g. De Lucia et al. 1994; Dimitrov et al. 1994; Khadake and Rao 1995; Ura et al. 1996; Fantz et al. 2001; Saeki et al. 2005), their biological function is also less clear. Single knock-outs might not always result in a physiological consequence (e.g. Lin et al. 2000; Fantz et al. 2001; Martianov et al. 2005; Tanaka et al. 2005). Nevertheless, ectopic expression, RNAi knock-down or antibody-mediated depletion of oocyte specific H1 variants did suggest a role in maintaining specialized chromatin structures (Funaya et al. 2018), and in regulating pluripotency of the early cells and facilitating nuclear reprogramming (e.g. Jullien et al. 2010; Hayakawa et al. 2012). How much of these results from *ex vivo* studies can be applied to the *in vivo* situation remains unclear.

*Drosophila melanogaster*, a genetically tractable organism with a well characterized developmental program, has contributed significantly to our understanding of the functions of H1 (for reviews see Hergeth and Schneider 2015; and Bayona-Feliu et al. 2016). Drosophila *his1* gene exists in a multi-copy gene cluster at the *histone* locus, similar to the situation in other higher eukaryotes, therefore genetic knock-out of *his1* has not been utilized for the dissection of H1 function even though recent advances in genome engineering has made it possible (Günesdogan et al. 2010; McKay et al. 2015; Zhang et al. 2019). Instead, prior functional studies of H1 relies on RNAi knock-down or overexpression approaches. It has been shown that H1 is important for the maintenance of heterochromatin stability and gene expression (Ni et al. 2006; Lu et al. 2009, 2013; Vujatovic et al. 2012), the regulation of DNA replication in polyploid cells (Andreyeva et al. 2017), the compaction of chromatin in endo-replicating cells (Corona et al. 2007; Lu et al. 2009; Siriaco et al. 2015), and stem cell maintenance in testis (Sun et al. 2015).

In 2013, the *bigH1* gene was discovered to encode the only H1 variant in the Drosophila genome (Pérez-Montero et al. 2013). Remarkably, *bigH1* has a germline specific pattern of expression making it a great model for the understanding of H1 variants in germline and early zygotic development. Indeed, both embryonic and male germline defects were reported for mutant animals (Pérez-Montero et al. 2013; Carbonell et al. 2017). However, the facts that BigH1 is loaded maternally, and that BigH1 is replaced by H1 before the activation of the zygotic genome, are conceptually inconsistent with the recessive zygotic lethality described for the *bigH1* mutant in the 2013 study. We generated new *bigH1* mutations and discovered that the loss of BigH1 has no discernable effect on development. We also provide evidence suggesting that maternal H1 can fully substitute BigH1 for early development.

## Materials and Methods

### Cas-9 mediated mutagenesis of *bigH1*

Mutations of *bigH1* were generated using a transgenic approach in which both the Cas9 protein (expressed from a *vasa* promoter) and gRNA (expressed from a *U6* promoter) were produced from transgenes inserted into the Drosophila genome (Port et al. 2014). The target gRNAs were designed with the online tool: http://tools.flycrispr.molbio.wisc.edu/targetFinder/. Target #1 is about 40bp from the annotated ATG start of *bigH1* and has the sequence of 5’-GGTTGAACGCAATGATGGCT**CGG** with the PAM sequence in bold. Target #2 is about 470bp from the annotated ATG start and has the sequence of 5’-GGCCGAAGCCAACGGCGAAG**TGG** with the PAM sequence in bold. Mutations were verified by genomic PCR and sequencing using DNA samples from homozygous mutant adults.

### Transgenic constructs

The rescuing *bigH1* transgene carries a 4.3 kb genomic fragment from the *bigH1* locus, containing the coding region plus an additional one kb each of the 5’ and 3’ regulatory regions. The method of recombineering was used to generate a *bigH1* gene tagged at the C-terminus with *gfp* as described (Zhang et al. 2014). Transgenes were introduced into the genome by standard phiC31-mediated insertion.

For live analyses of H1 and H2B distribution in germline cells, a transgene carrying a single 5kb unit of the *histone* cluster (McKay et al. 2015) was first constructed. For inserting *gfp* or *mcherry* tags into *his1* or *his2b* coding regions, the method of recombineering was employed as described (Gao et al. 2009; Zhang et al. 2014). The coding regions of *his1* and *his2b* were swapped using recombineering. These constructs with various configurations of *his1* and *his2b* expression (Figure 4A) were introduced into the genome by phi-C31 mediated insertion.

### Immunostaining and Western blot analyses

Antibodies against BigH1 (from mice) and H1 (from guinea pigs) were raised against full length recombinant proteins as antigens purified from bacteria. The specificity of anti-BigH1 was shown in the Western blots in Figure 1 and immunostaining experiments in Figure 2. The specificity of anti-H1 was shown in Western blots in Figure S5.

**Figure 1.**
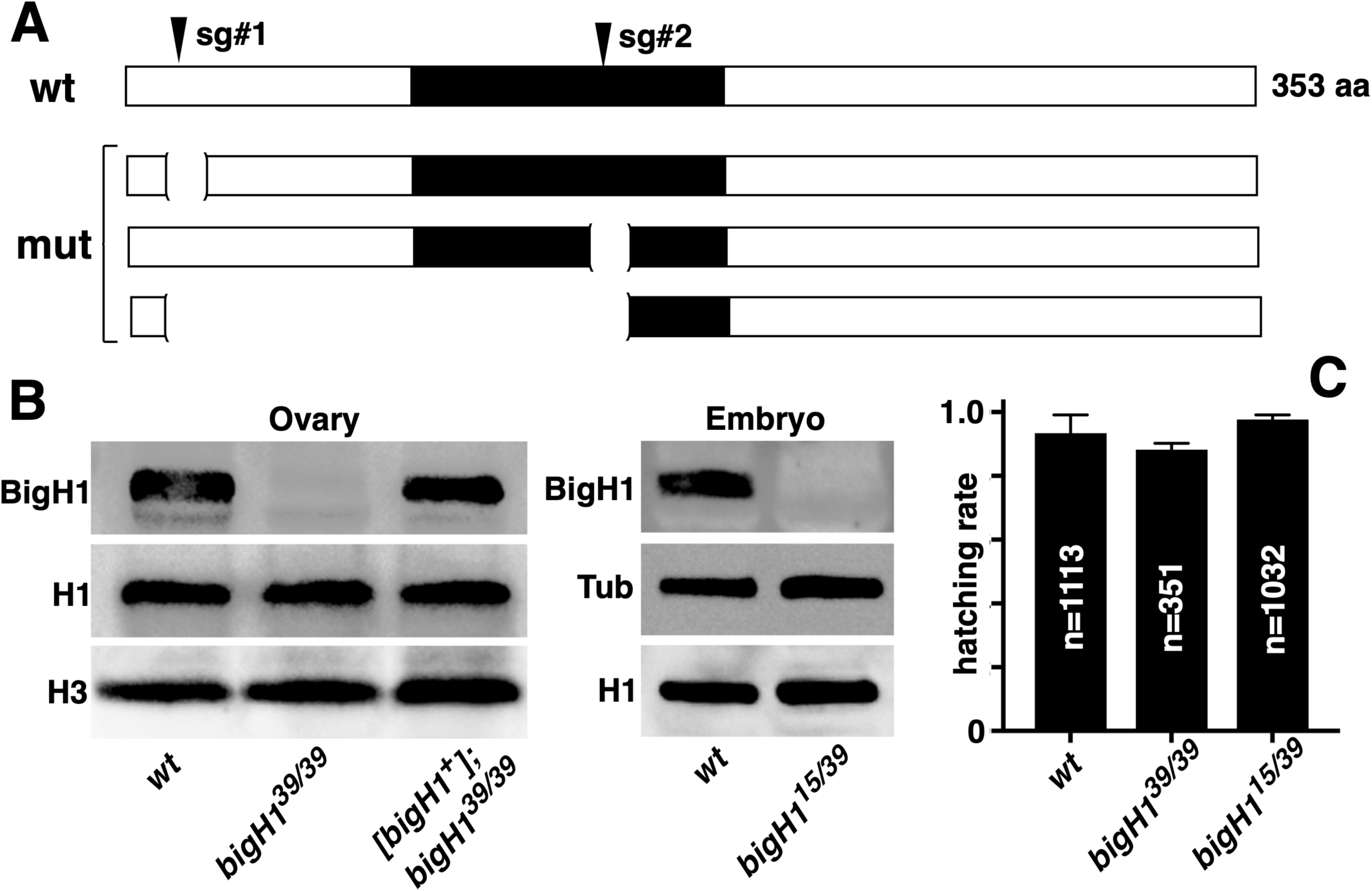
Null mutations of *bigH1* support normal development. **A**. New *bigH1* mutations. The diagram at the top depicts the coding region for the 353 amino acid long BigH1 protein with the conserved globular domain shown in black. The approximate positions are indicated for the two guide RNAs (sgRNA) used in Cas9-mediated mutagenesis, as well as the deleted regions in three classes of mutations. **B**. Western blot analyses showing that *bigH1* mutants lack the BigH1 protein. Ovarian and embryonic (0-15min) extracts were used. Flies with the genotype *[bigH1*^*+*^*]; bigH1*^*39/39*^ carried a *bigH1* wild-type transgene (rescued flies). Mutant extracts have normal level of H1. Histone H3 and tubulin were used as loading controls. **C**. New *bigH1* mutations support normal hatching rates of embryos.

**Figure 2.**
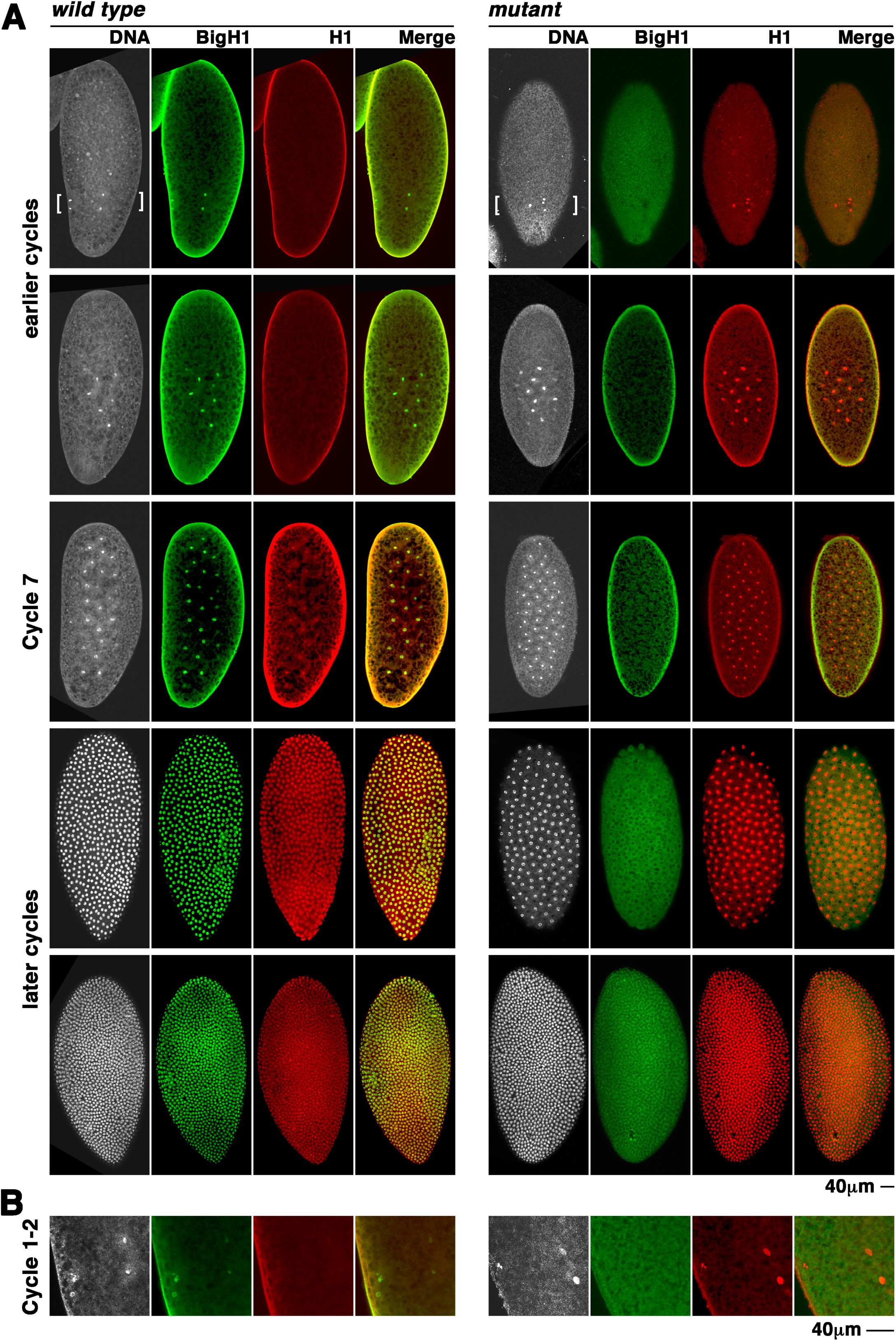
Compensatory loading of H1 in the absence of BigH1. Syncytial embryos of wild type and *bigH1*^*15/39*^ (*mutant*) were stained with DAPI (in white), anti-BigH1 (in green) and anti-H1 (in red) antibodies. The channel for each dye is shown separately as well as the merged image of BigH1 and H1 signals. In all pictures, the posterior of the embryo is up. **A**. Pictures of whole embryos displayed according to their developmental stages (in cycles earlier than cycle 7, cycle 7, and cycles later than 7). In very early cycles, the areas where DAPI-stained nuclei can be seen are marked with white brackets. Cycle 7 is when chromatin H1 signals was first detected in wild type embryos. **B**. Higher magnification of embryos highlighting that H1 can be detected as early as cycle 1-2 in *bigH1* mutants.

Embryos staining was performed by standard protocols. Briefly, 0-2h embryos were collected and fixed in a 1:1 mixture of 4% formaldehyde and heptane for 20 minutes. Fixed embryos were stained with the following primary antibodies: mouse anti-bigH1 (1:500) and guinea pig anti-H1 (1:500) followed by secondary antibodies at 1:500. For Western blotting, protein extracts were prepared by homogenizing embryos or ovaries in lysis buffer (50 mM Tris-HCl pH 8.0, 100 mM NaCl, 0.1% Triton X-100, 1X protease inhibitor cocktail (Roche)). After lysis, 4X SDS-PAGE loading Buffer was added to the extract, and samples were boiled. The primary antibodies were used at a dilution of 1:5,000.

### Partial micrococcal nuclease (MNase) digestion assay

The assay was performed as described by Xu et al. (2016). Briefly, 0-2h embryos were homogenized in 200 μl of Buffer A. Nuclei were pelleted, washed twice with Buffer B, and resuspended in 200 μl of Buffer B. Nuclei were digested with MNase (from NEB) per manufacturer’s instruction for 10 min at 37°C. The digestions were stopped with EDTA, RNA was degraded with RNase A for 20 min at 37 °C, and samples were treated with Proteinase K followed by a phenol-chloroform extraction. Precipitated DNA was dissolved in TE and loaded on a 1.5% agarose gel in 1x TAE. The gel was stained with EtBr after electrophoresis.

The estimation of nucleosomal spacing was performed essentially as described by Potdar et al. (2018). Briefly, a standard curve was generated using the ladder of molecule markers. The size of DNA fragments carrying different numbers of nucleosome particles was calculated and plotted. The slope of the line represents the estimated nucleosome distance in base pairs.

### Live imaging

Live analyses of embryos were performed as described (Fasulo and Sullivan 2014). Briefly, embryos expressing green fluorescent H2Av proteins were collected every hour, dechorionated by hand, aligned and glued to glass coverslips, and covered with halocarbon oil before imaging. Embryonic development was recorded for 1-1.5h with 40s intervals using an Olympus confocal microscope under a 20x objective.

Live analyses of HP1 occupancy at the 359 satellites were conducted as described (Yuan and O’Farrell 2016). Briefly, GFP-HP1 and TALE-mCherry recombinant proteins were purified from bacteria and injected into syncytial embryos, which were imaged to record the movement of both GFP and mCherry tagged proteins. The accumulation of GFP-HP1 on TALE-mCherry marked 359 repeats was quantified as described in Yuan and O’Farrell (2016), particularly in their Figure S3.

### qPCR

Expression analyses of typical zygotic genes in embryos were performed as described with the same primer sets for qPCR (Pérez-Montero et al. 2013).

## Results and Discussion

### Null mutations of *bigH1* support viability and fertility

Previously, the *bigH1* gene was identified to encode a variant of histone H1 in Drosophila (Pérez-Montero et al. 2013). It was also discovered that *bigH1* is exclusively expressed in the germline, that the BigH1 protein is maternally loaded into the egg, and that chromatin-bound BigH1 is gradually replaced by somatic H1 as fertilized eggs develop into zygotically activated embryos. The *bigH1*^*100*^ allele was reported as a recessive zygotic lethal mutation at the embryonic stage. This is rather surprising considering that homozygous mutant embryos had heterozygous mothers that would have deposited abundant BigH1 protein into the egg. It would have been more consistent with BigH1’s distribution pattern if *bigH1* mutants had displayed maternal effects on embryonic development. We decided to investigate the possible cause for this discrepancy using new *bigH1* alleles that we generated by CRISPR/Cas9-mediated mutagenesis.

As described in Materials and Methods and shown in Figure 1A, we used guide RNAs (gRNA) to target two regions of the *bigH1* gene, one just downstream of the annotated ATG codon and one at a region predicted to encode the conserved globular domain (Pérez-Montero et al. 2013). We recovered multiple frameshift mutations at both regions. By the simultaneous usage of both gRNAs, we were able to recover multiple deletion mutations of *bigH1* (Figure 1A and Table S1). We decided to carry further analyses with deletion mutations as well as frameshift mutations at the globular domain as they are most certainly null or very strong loss-of-function mutations. As a precaution, we used these two different kinds of mutations as trans-heterozygous animals in many of our analyses so to minimize the potential effect of second site mutations in the background.

Western blot with an anti-BigH1 antibody confirms the absence of BigH1 protein in ovarian and embryonic extracts taken from mutant animals (Figure 1B). For all *bigH1* mutant combinations tested, homozygous animals were recovered at Mendelian ratios from heterozygous crosses (N>5000). As shown in Figure 1C, *bigH1* mutant males and females have normal level of fertility, and homozygous mutant stocks have been stably maintained for all alleles. We conclude that *bigH1* is not essential for animal viability.

In all of our subsequent analyses of BigH1’s function in embryonic development, we used cleavage stage embryos produced by homozygous mutant mothers. We called these *bigH1*-mutant embryos because they lack BigH1 proteins that would have been deposited by the mother normally.

### Compensatory loading of embryonic H1 in the absence of BigH1

Because animals with reduced levels of the somatic H1 are inviable (Lu et al. 2009; Vujatovic et al. 2012), we had expected that *bigH1* mutations would have caused a strong maternal effect, possibly lethality, in that embryos produced by *bigH1* mutant mothers would not have survived. Therefore, our results that *bigH1*-mutant embryos survive at a normal rate (Figure 1C) raise the question of what fulfills the function of H1 in these embryos. As it is known that H1 is maternal deposited, even though it is scarcely if at all present on chromatin in the early cell cycles (Becker and Wu 1992; Ner and Travers 1994), we investigated the possibility that chromatin loading of H1 is upregulated in the absence of BigH1 therefore supplying the “H1” function.

Using Western blot on embryos collected every 15 minutes, we demonstrate the presence of H1 in early embryos (Figure 1B). Interestingly, these H1 molecules are mostly if not at all absent from chromatin in immunostaining experiments (Figure 2). Consistent with results from an early study (Ner and Travers 1994), we detected signals of chromatin bound H1 starting at Cycle 7 in wild type embryos (Figure 2A). Remarkably in *bigH1* mutant embryos, H1 is present as early as the first cycle (Figures 2A and B). More specifically, chromosomal signals of H1 were observed in none of the 59 wild type but all of the 61 mutant embryos that were in a cycle earlier than 7. In contrast, all embryos at or after cycle 7 had H1 signals on chromatin regardless of their maternal genotype (n=47 for wild type and 76 for mutant). These results support our previous proposition that H1 loading becomes earlier and possibly stronger in *bigH1*-mutant embryos.

Our immunostaining results were not meant to provide an accurate estimation of how much of the BigH1 occupancy on chromatin was replaced by that of H1. We surmise nevertheless that the degree of replacement must have been sufficient to be detected by our staining method and to provide H1 function to support normal development. Importantly, this “ectopic” H1 loading in the mutants can be “rescued” by the introduction of a wild type *bigH1* transgene. As shown in Figure S1 in Supplemental Materials, these “rescued” embryos display an H1 immuno-localization pattern identical to that of the wild type embryos.

### Replacement of BigH1 with maternal H1 does not alter chromatin structures or cell cycle progression

As an assessment of the general chromatin structure in *bigH1* mutant embryos, we employed the classical Micrococcal Nuclease (MNase) assay in which the native chromatin is partially digested with MNase to characterize nucleosome occupancy (Becker and Wu 1992). As shown in Figures 3A, the estimated distance between nucleosomes is not changed upon loss of BigH1, a conclusion not consistent with the one reached in the 2013 study. Given that we showed compensatory loading of H1 in the mutant, this is not an unexpected result since H1/BigH1 occupies the linker region between nucleosomes even though the two H1s might differ in sequence and biochemical properties. H1 has been shown to facilitate chromatin compaction (reviewed in Zhou and Bai 2019), whether higher orders of chromatin structure are altered upon H1’s replacing BigH1 awaits further investigation.

**Figure 3.**
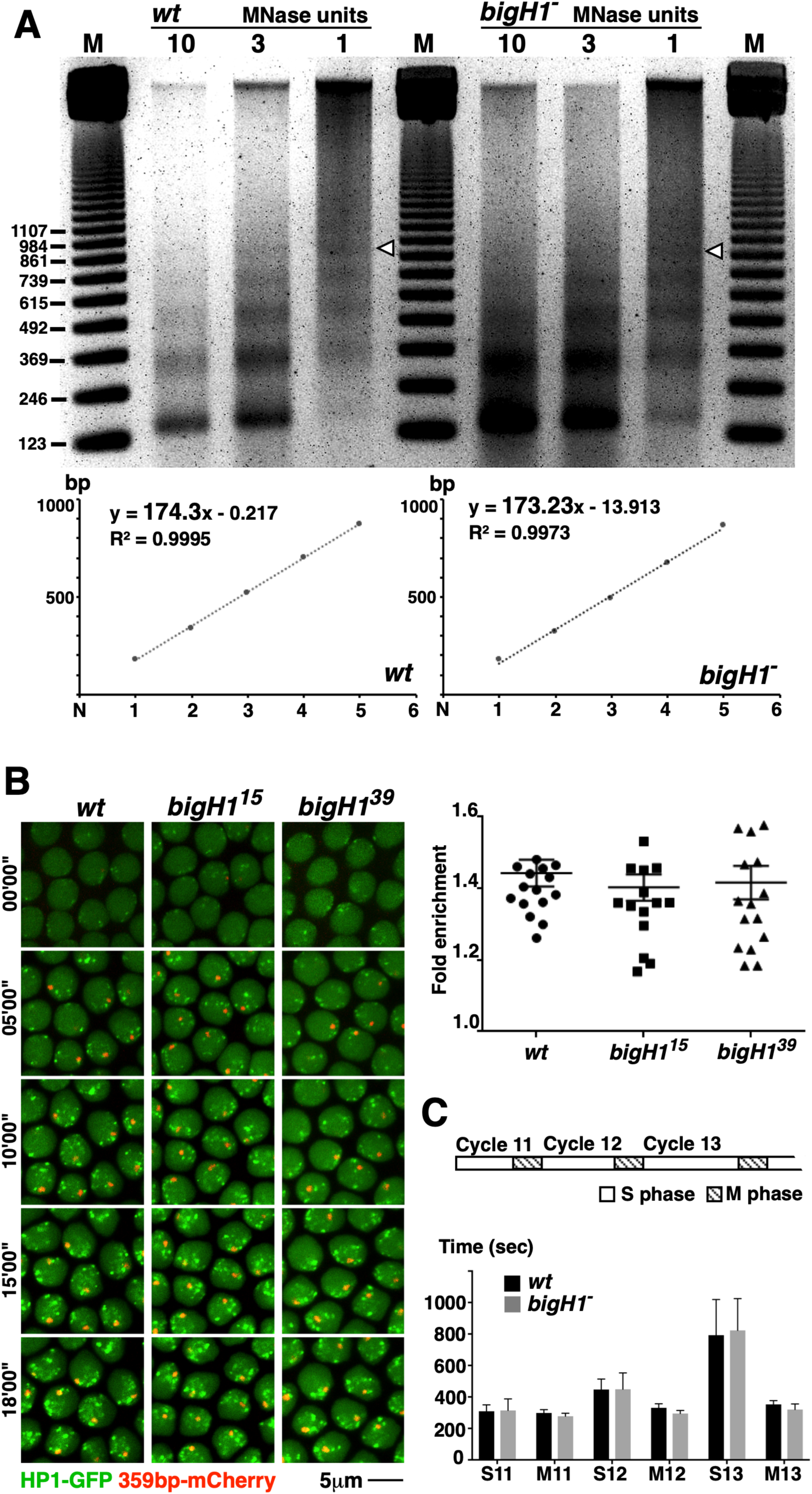
Normal chromatin structures and cell cycle progression in *bigH1*-mutant embryos. **A**. The MNase digestion assay for estimating nucleosomal spacing. Nuclei of syncytial embryos from *wt* and *bigH1*^*39/39*^ (*bigH1*^*-*^) females were digested with decreasing amount of MNase (left to right) and genomic DNA along with molecular markers (sizes in base pairs) were ran on a gel and stained. The approximate position of the DNA band corresponding to the penta-nucleosomal fragment is marked with a white arrowhead. The molecular sizes of different nucleosomal fragments were estimated and plotted with N denoting the number of nucleosomes (lower panels). The slope, which represents the space between nucleosomes, were estimated to be 174bp for *wt* and 173bp for the mutant samples. **B**. The dynamics of HP1 localization to heterochromatic satellites. Syncytial embryos (n>3 for each genotype) with the indicated genotypes (top) were injected with purified GFP-HP1 and TALE-mCherry proteins, which binds to the 359 repeats. The interphase of Cycle 14 was imaged and shown (left panel), which allows a visual inspection of the dynamics of HP1 and 359 co-localization. The extent of GFP signals colocalizing with mCherry signals was estimated for over 16 individual nuclei for each genotype. The “Fold enrichment” was calculated for each nucleus and plotted (right panel). No significant difference was observed between wildtype and *bigH1*^*15*^ embryos (unpaired t-test, P=0.7063), or between wildtype and *bigH1*^*39*^ embryos (unpaired t-test, P=0.9295). Error bars represent the SD. **C**. Cell cycle durations by live analyses. At the top is an illustration of a typical cell cycle profile for the last three syncytial cycles (11, 12, 13) showing the S and M phases separately. At the bottom is the measured lengths of S and M phases for each cycle. For wildtype, n=11. For the mutant (*bigH1*^*15/39*^), n=14. For a scatter plot representation of the same data set see Figure S2, which includes statistical analyses.

**Figure 4.**
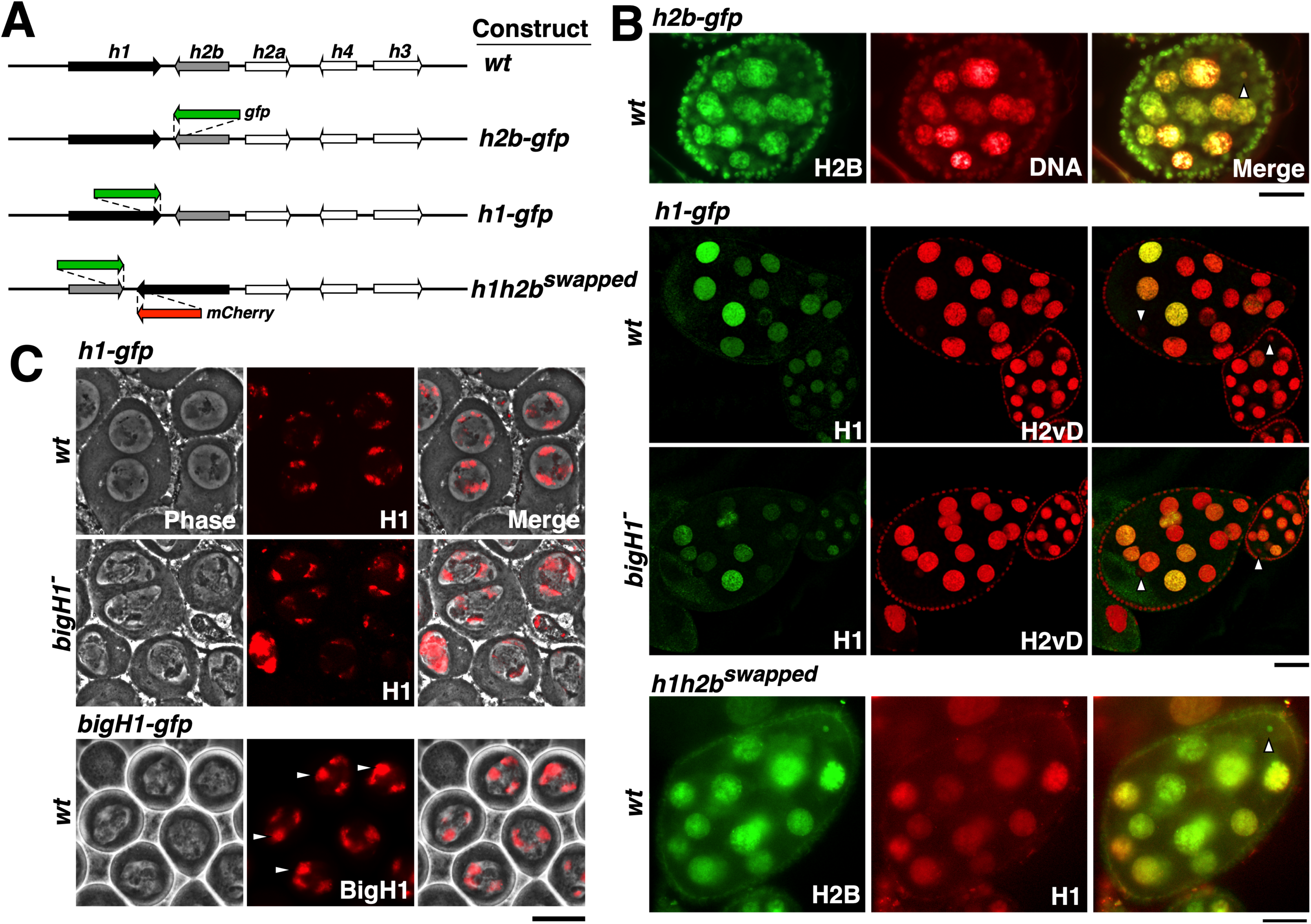
Germline distributions of H1 and BigH1 histones. **A**. Transgenic constructs expressing fluorescently tagged histones. At the top is the 5kb wildtype *histone* unit with coding regions of the five histone genes shown as rectangles with arrows pointing to the direction of transcription. Gene name abbreviations: *h1*=*his1, h2a*=*his2a, h2b*=*his2b, h3*=*his3, h4*=*his4*. In the “*h2b-gfp*” construct, the coding region of *gfp* was inserted at the C-terminus of *h2b*, and it was inserted at the C-terminus of *h1* in the construct “*h1-gfp*”. In the “*h1h2b*^*swapped*^” construct, the positions of the coding regions of *h1* (tagged with *mCherry*) and *h2b* (tagged with *gfp*) were exchanged. **B**. Histone localization in live egg chambers. The names of the construct used to produce each panel group are listed at the top. The top panel shows an egg chamber with H2B-GFP (in green) and DNA (stained with Hoechst, in red) channels shown separately. Also shown is the merged image in which the position of the oocyte nucleus is marked with a white arrowhead. In the middle two panels, H1-GFP (in green) and H2AvD-RFP (in red) signals are shown in addition to the merged images in which the oocyte nuclei are marked with white arrowheads. In both wildtype (*wt*) and *bigH1*^*15/39*^ (*bigH1*^*-*^) mutant egg chambers, H1 is undetectable in the nuclei of the oocytes. In the bottom panel, H2B-GFP (in green) under the control of the *h1* regulatory elements and H1-mCherry (in red) under those of the *h2b* are shown separately in addition to the merged image. The arrowhead marks the nucleus of the oocyte, which displays H2B but not H1 signals. More than 100 egg chambers were observed for every genotype. **C**. Histone localization in live testis. The names of the construct used to produce each panel group are listed at the top. Phase contrast images showing pre-meiotic spermatocytes with condensed chromosomes displaying H1-GFP or BigH1-GFP signals (both in red). H1 localization in wildtype (*wt*) and *bigH1*^*15/39*^ (*bigH1*^*-*^) mutant testes are similar. In particular, BigH1-GFP forms a prominent and large domain (marked with white arrowheads), possibly representing the *rDNA* loci on the sex bivalent. In the absence of BigH1, H1 does not form a similar domain indicative of the lack of extra loading of H1.

H1 has been extensively implicated in heterochromatin formation in Drosophila somatic cells (Vujatovic et al. 2012; Lu et al. 2013), we investigated whether BigH1 regulates this feature of chromatin structure during early divisions. Using live imaging we monitored the accumulation of Heterochromatin Protein 1 (HP1) over a large block of repetitive sequence (the 359 satellite) in heterochromatin of the *X* chromosome (Yuan and O’Farrell 2016). This assay was recently employed to show that the chromatin regulator Eggless/SetDB1 governs heterochromatin formation during embryonic development (Seller et al. 2019). As shown in Figure 3B, neither the timing nor the level of HP1 accumulation at the 359 satellite differ between wildtype and *bigH1*-mutant embryos.

Although *bigH1* embryos hatched at normal rate, there might be subtle defect in cell cycle progression that would not be detectable in our hatching measurement. We set out to measure the length of the cell cycles by live imaging of embryonic nucleic marked with fluorescently tagged histones (see Materials and Methods). As nuclei in early embryos arrive at the apex during the 10th syncytial cycles, we were able to capture, to completion, the last three syncytial cycles (Cycles 11, 12 and 13) before zygotic activation. Representative images of both wild-type and *bigH1* mutant embryos are shown in Figure S2A in Supplemental Materials. As summarized in Figure 3C and shown in more details in Figures S2B and S2C, early cell cycles in the mutant progress at similar rates as those in wild-type embryos, regardless of whether the total cycle length or the length of the interphase were measured. However, we observed a statistically significant shortening of mitotic division for each of the last three cycles in the mutant. The underlying cause for this difference is not known, but it nevertheless does not affect the viability of the embryos. We conclude therefore that both chromatin structures in bulk and cell cycle progression in general are normal in BigH1-deficent embryos.

### Compensatory replacement of BigH1 with H1 might be specific for embryonic cells

We are interested in whether this wholesale BigH1 to H1 replacement is specific to cells in early embryos as these cells undergo rapid division and dynamic chromatin remodeling. We studied the germline where BigH1 is specifically present, and focused on germline cell type(s) in which the two H1 proteins display different patterns.

In the ovary, BigH1 is prominently present in the oocyte nucleus (Pérez-Montero et al. 2013; and Figure S3 in Supplemental Materials). To monitor H1 localization, we constructed a single *histone* gene cluster in which a 5kb genomic fragment contains all five canonical histones under the control of their own regulatory elements (Figure 4A). We then tagged *his1* with *gfp* in the transgenic construct. H2B was separately tagged with GFP and used as a representative for the nucleosomal histones. When we introduced these constructs into the genome, we observed robust H2B-GFP signals in the ovaries (Figure 4B). In particular, H2B-GFP is present in the nucleus of the oocyte. In comparison, H1-GFP is not detectable in the oocyte nucleus even though it is present in nuclei of the large polyploid nurse cells (Figure 4B). We were interested in whether the presence of BigH1 causes H1 being undetectable in the oocyte nucleus, as our “compensatory replacement” model predicts. When the *his1-gfp* construct was introduced into *bigH1* mutant ovaries, H1-GFP is again undetectable in the oocyte nucleus (Figure 4B), in contrast with a robust presence of H2B-GFP in the same cells. Therefore, the absence of BigH1 does not lead to appreciable chromatin loading of H1 in the oocyte nucleus. However, due to the relatively low level of H1 in comparison with H2B, we cannot rule out a small increase of H1 in *bigH1*-mutant oocytes.

It was shown previously that expression of *his1* relies on TRF2 rather than the TBP transcription factor that is used for the other *histone* genes, which results in different distribution patterns during the S phase between H1 and the other histones (Isogai et al. 2007). We wished to investigate whether this special mode of *his1* expression causes the difference in histone distributions inside the oocyte. We constructed a third construct in which the coding regions for H1 and H2B were swapped so that *his1* expression is now under the control of the *his2b* regulatory mode and *vice versa* (Figure 4A). As shown in Figures 4B, this promoter-swapping did not result in discernable changes in H2B or H1 localization, thus ruling out that difference in transcriptional regulation is responsible for the difference in protein distribution.

The lack of a H1 uploading in the absence of BigH1 was similarly observed in the testis. As shown in Figure 4C, BigH1-GFP is localized on condensed chromosomes in spermatocytes, with a prominent presence at a domain that likely corresponds to the *rDNA* loci on the sex bivalent. H1-GFP, on the other hand, is not prominently localized to the same domain despite its presence on condensed chromosomes. More importantly, the loss of BigH1 does not alter the pattern of H1-GFP distribution (Figure 4C).

These results suggest that the compensatory loading of H1 onto BigH1 deprived chromatin occurs to a much larger extent during early embryonic than germline development.

### Concluding remarks

Our investigations into the function of BigH1 have produced results that are inconsistent with those from two prior reports. Pérez-Montero et al. (2013) concluded that even in the presence of the maternal pool of BigH1, *bigH1* homozygous embryos experience changes in chromatin structure, premature activation of zygotic transcription, and ultimately death. Carbonell et al. 2017 concluded that reducing BigH1 in testis results in a severe disruption of male germline development. In contrast, our *bigH1* null mutants have normal viability and fertility. At the cellular level, *bigH1* mutant embryos have normal cell cycle progression, normal nucleosomal spacing, normal heterochromatin establishment and no premature activation of typical zygotic genes (Figure S4). The exact cause for these discrepancies is not known although we suspect that it has to do with the ways used to disrupt BigH1 function in the prior studies (a single allele (*bigH1*^*100*^) with nucleotide changes in the non-coding region of *bigH1* in the 2013 study and germline specific RNAi in the latter). Our results instead demonstrate a remarkable compensatory activity that must have been activated upon the loss of an important chromatin protein. Such a phenomenon has been previously shown also in Drosophila in which overexpression of BigH1 and the BEN domain protein Elba2 could partially substitute for the loss of H1 (Xu et al. 2016). In addition, the HMG-D protein has been implicated as one of the proteins that might share similar function as H1 in Drosophila (Ner and Travers 1994; although also see Nalabothula et al. 2014). Furthermore, loss of the mouse testis specific variant H1t led to an increased presence of somatic H1 on testis chromatin (Fantz et al. 2001). Therefore, the specific function of BigH1 that distinguishes it from H1, which must have given its existence an evolutionary advantage, awaits future investigation.

## Supporting information

manuscript

## Acknowledgements

We thank Dr. Jinyun Ji for her assistance in constructing the original *histone* construct. We thank Mr. Qianyi Zhang for his assistance in characterizing some of the *bigH1* mutations. This work was supported by the National Key R&D Program of China (2018YFA0107000) to YSR, by the National Natural Science Foundation of China (31771589) and the Department of Science & Technology of Hunan Province, China (2018DK2015, 2017XK2011,2017RS3013) to KY, and by the Intramural research program of the National Cancer Institute, USA to YB.

## Author contributions

Conceptualization, KKL and YSR; Methodology, KKL, YSR; Formal Analysis, KKL, KY and YSR; Investigation, KKL, DH, FC, RL, BZ, KY, and YSR; Resources, BZ and YB; Writing-Original Draft, YSR; Writing-Review & Editing, KKL, KY, BZ, YB and YSR; Supervision, YB, KY and YSR; Funding Acquisition, YB, KY and YSR.

## Declaration of Interests

The authors declare no competing interests.

