## Supplementary material for "Compensatory replacement of the BigH1 variant histone by canonical H1 supports normal embryonic development in Drosophila": manuscript

**Figure Legends for Supplemental Materials**

**Figure S1. Re-supplying BigH1 restores restrictive loading of H1 to embryonic chromatin in *bigH1* mutants**

Syncytial embryos shown were from *bigH1*$^{15/39}$ mutant females that also carry a wildtype *bigH1* transgene (rescued individuals). They were stained with DAPI for DNA (in white), anti-BigH1 (in green) and anti-H1 (in red) antibodies. The channel for each dye is shown separately as well as the merged image of BigH1 and H1 channels. In all pictures, the posterior of the embryo is to the right. Embryos are displayed according to their developmental stages (in cycles earlier than cycle 7, cycle 7, and cycles later than 7). In the early embryo, three BigH1-positive clusters can be seen while only two of them have corresponding DAPI signals that are above the background (black arrows). The third cluster, marked by an arrowhead, is not detectable beyond the background level. Cycle 7 is when chromatin H1 signals were detected in these rescued embryos similar to the wild type situation. For embryos earlier than cycle 7, n=115, and n=75 for embryos at cycles 7 or later.

**Figure S2. Representative images of nuclear morphology from live imaging of syncytial divisions and data analyses of cell cycle lengths**

**A**. Embryos from wild type (*wt*) and *bigH1*$^{15/39}$ (*bigH1*$^{-}$) females are imaged. Green fluorescence from tagged histones is shown in greyscale pictures. In the *wt* embryo, telophase was recorded at 00:40 for cycle 10 and at 10:40 for cycle 11, giving rise to a cell cycle length of about 10 minutes. In the mutant embryo, telophase was recorded at 02:00 for cycle 10 and at 11:20 for cycle 11, giving rise to a cell cycle length of about 9 minutes and 20 seconds. **B**. Scatter plots for M phase duration (left) and total cell cycle duration (right) using the same data set as those in Figure 3C. **C**. Data analyses of cell cycle measurements. The mean durations (in seconds) of the S phase, the M phase and the entire cell cycle for syncytial cycles 11-13 are tabulated for wild type (*wt*) and *bigH1*$^{15/39}$ (*bigH1*$^{-}$) embryos, along with the SD. The differences in length were calculated by subtracting the mean of the mutant from the mean of the *wt* samples. The P values from unpaired t-tests comparing the two samples are listed with statistically significant comparisons ($p<0.05$) shown in bold.

**Figure S3. BigH1 localization in the ovary**

Wildtype ovaries were stained with anti-BigH1 (in green) and DAPI (in red). BigH1 is prominently present in the oocyte nuclei, which are labelled with white arrowheads.

**Figure S4. Loss of BigH1 does not activate typical zygotic genes**

qPCR analyses were performed precisely as described in Pérez-

Montero et al. 2013. The genes analyzed and primer sets are identical to those in the prior study. Data analyses are represented below the chart. No pairwise comparison between wildtype and the mutant shows significant difference in expression level.

**Figure S5. The specificity of anti-H1 antibodies**

Western blot analyses on extracts taken from larval salivary glands. The “control” larvae carried an *actin-gal4* transgene, and the “rnai” larvae carried *actin-gal4* and a *UAS* construct expressing siRNAs against the Drosophila *his1* gene, which led to a reduction of H1 when compared with the Tubulin loading control.

Table S1. Sequences of *bigH1* mutant alleles

| | Allele[1] | Sequences[2, 3, 4] | | |
|---|---|---|---|---|
| | *wt* | CGCAATGATGGCT**CGG**ATGA | (411nt) | GGCCGAAGCCAACGGCGAAG**TGG**TGATGGTTAAGCGA |
|  | 9-11(△8nt) | CGCAATG-----------------GATGA | (411nt) | GGCCGAAGCCAACGGCGAAGTGG |
| Target #1 | 3-11(△2nt) | CGCAATGA----GCTCGGATGA | (411nt) | GGCCGAAGCCAACGGCGAAGTGG |
| | 4-12(△1nt) | CGCAATGATG--CTCGGATGA | (411nt) | GGCCGAAGCCAACGGCGAAGTGG |
|  | 15(△5nt) | CGCAATGATGGCTCGGATGA | (411nt) | GGCCGAAGCCA-----------GAAGTGG |
| Target #2 | 41(▽8nt) | CGCAATGATGGCTCGGATGA | (411nt) | GGCCGAAGCCAACGGtggttggcCGAAGTGG |
|  | 21(▽5nt△9nt) | CGCAATGATGGCTCGGATGA | (411nt) | GGCCGAAGCCAccgct-------------TGG |
|  | 39(△440nt) | CGCAATGATGG-------------------- | (411nt) | --------------------------------------------TGG |
| Target 1+2 | 37(▽2nt△437nt) | CGAATGATGat--------------------- | (411nt) | ------------------------------------GAAGTGG |
|  | 26(△449nt) | CGCAATGATGG-------------------- | (411nt) | ------------------------------------------------------------ TTAAGCGA |

[1]: The names of mutant allele are given followed by the nature of the mutation in parentheses (△: deletion of nucleotides; ▽: insertion of nucleotides).

[2]: The PAM sequences of the target gRNAs are given in bold in the wildtype (*wt*) sequence.

[3]: There are 411nt between the two gRNA targets.

[4]: The deleted sequences are marked with "-" and the inserted sequences were given as small cap letters.

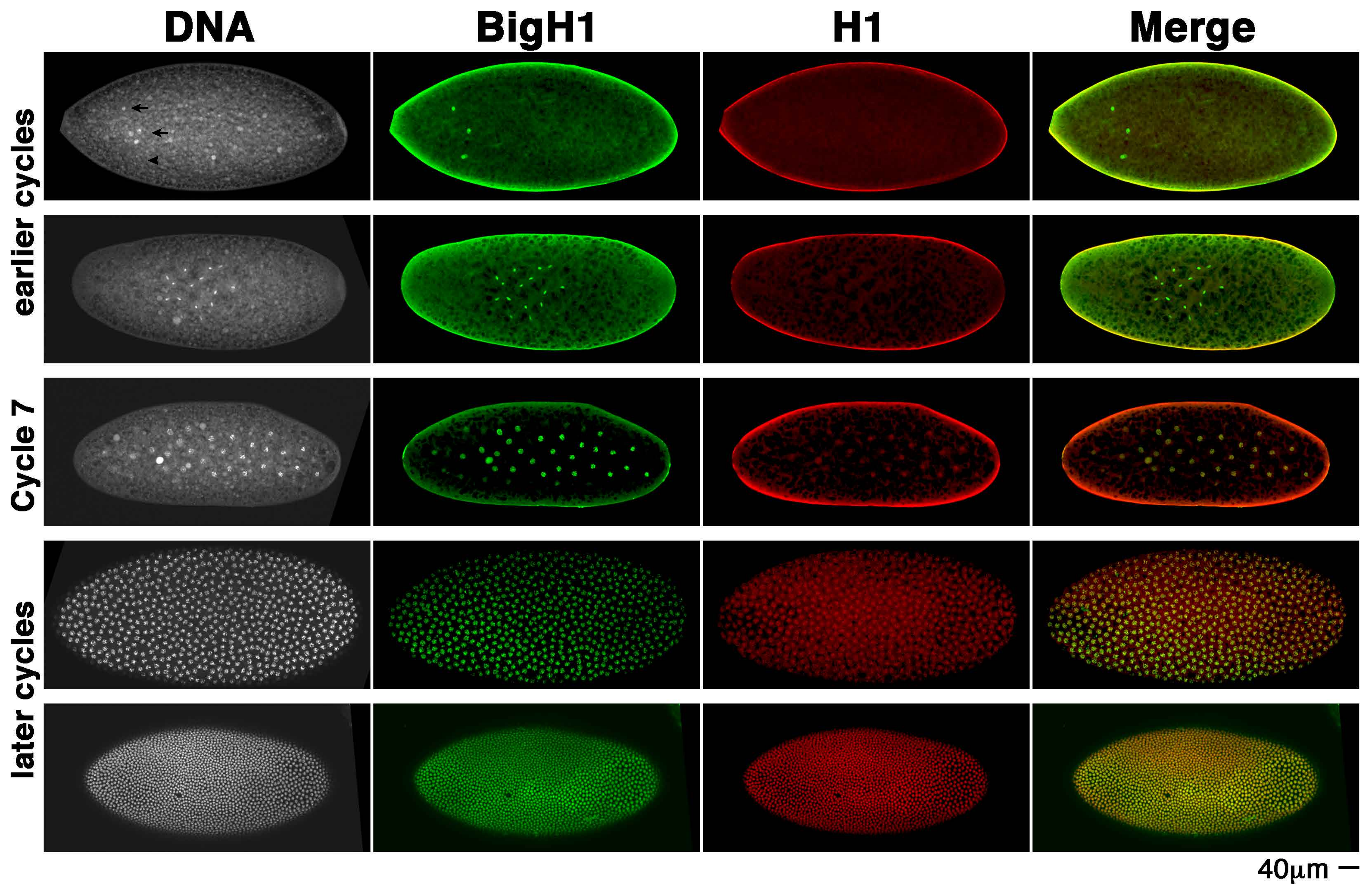
DNA
BigH1
H1
Merge
earlier cycles
Cycle 7
later cycles
40μm

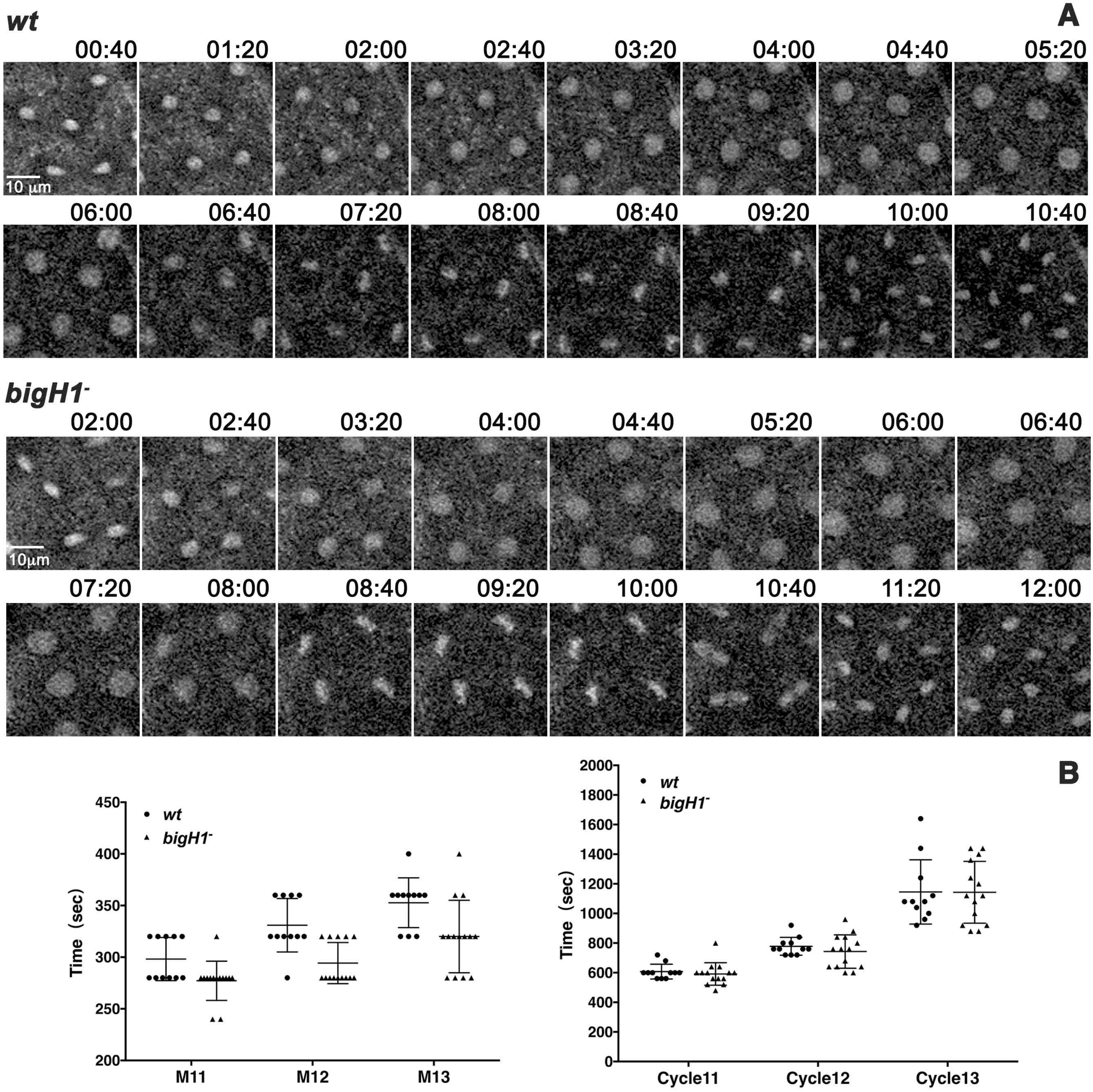


C

| | | S11 | M11 | S12 | M12 | S13 | M13 | Cycle11 | Cycle12 | Cycle13 |
|---|---|---|---|---|---|---|---|---|---|---|
| *wt* (N=11) | Mean | 309.1 | 298.2 | 447.3 | 330.9 | 792.7 | 352.7 | 607.3 | 778.2 | 1145.5 |
| | SD | 40.4 | 20.9 | 66.5 | 25.9 | 226.1 | 24.1 | 50.0 | 60.3 | 217.1 |
| *bigH1*⁻ (N=14) | Mean | 314.3 | 277.1 | 448.6 | 294.3 | 822.9 | 320.0 | 591.4 | 742.9 | 1142.9 |
| | SD | 73.3 | 19.0 | 104.3 | 19.9 | 202.0 | 35.1 | 75.5 | 112.8 | 208.6 |
| Difference | | -5.2 | 21.0 | -1.3 | 36.6 | -30.1 | 32.7 | 15.8 | 35.3 | 2.6 |
| P value | | 0.8351 | **0.0149** | 0.9717 | **0.0006** | 0.7285 | **0.0147** | 0.5552 | 0.3591 | 0.976 |

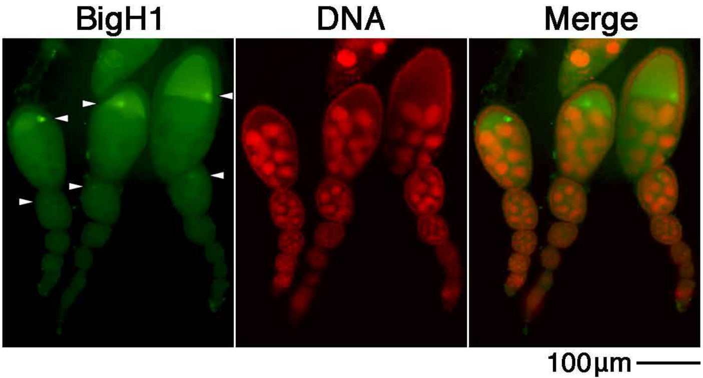
BigH1
DNA
Merge
100µm

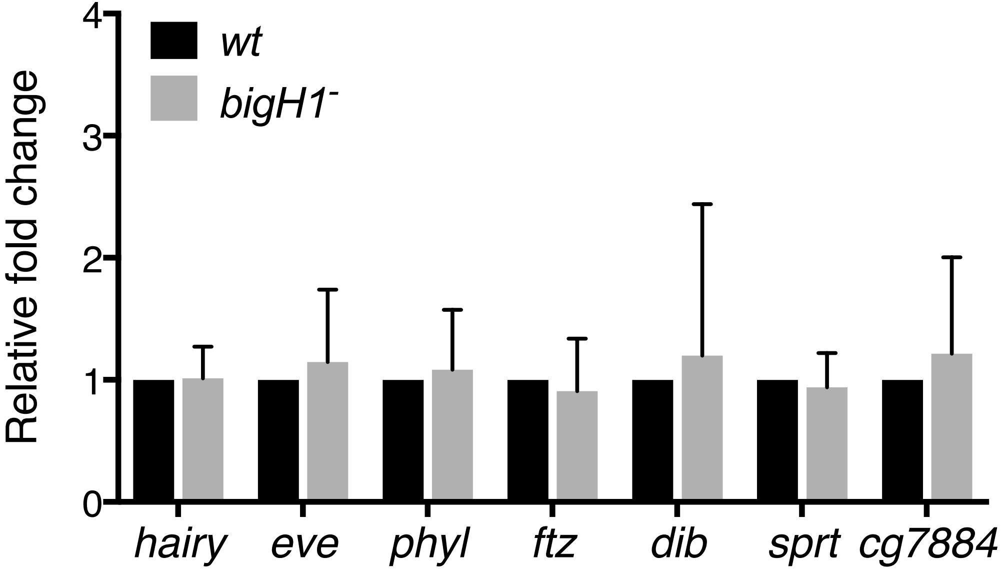


| | P value | *wt* | *bigH1*$^{-}$ | Difference | SE of difference | t ratio | df |
|---|---|---|---|---|---|---|---|
| *hairy* | 0.8894 | 1.00 | 1.02 | -0.02 | 0.1052 | 0.1426 | 10 |
| *eve* | 0.5586 | 1.00 | 1.15 | -0.15 | 0.2424 | 0.6051 | 10 |
| *phyl* | 0.6569 | 1.00 | 1.08 | -0.08 | 0.1851 | 0.4554 | 12 |
| *ftz* | 0.5836 | 1.00 | 0.91 | 0.09 | 0.1623 | 0.5633 | 12 |
| *dib* | 0.6770 | 1.00 | 1.20 | -0.20 | 0.4685 | 0.4269 | 12 |
| *sprt* | 0.5884 | 1.00 | 0.94 | 0.06 | 0.1053 | 0.5560 | 12 |
| *cg7884* | 0.5201 | 1.00 | 1.22 | -0.22 | 0.3225 | 0.6667 | 10 |

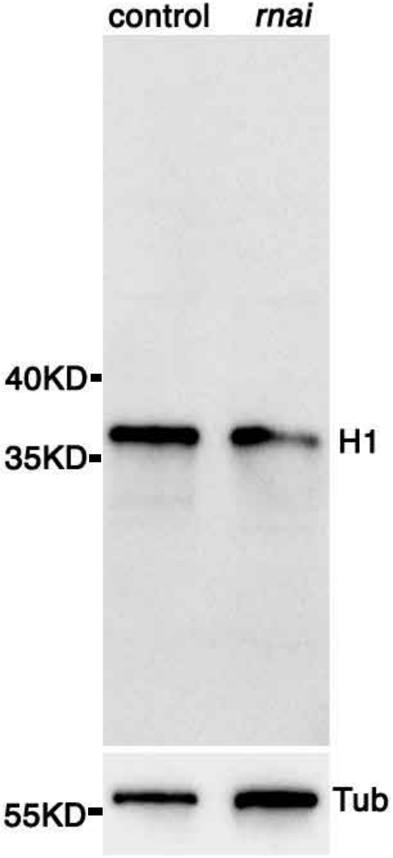
control
rnai
40KD
35KD
H1
55KD
Tub
